# Distribution, and abundances of Peleng Tarsier (*Tarsius pelengensis*), in Banggai Island group, Indonesia

**DOI:** 10.1101/2020.09.21.305888

**Authors:** Fakhri Naufal Syahrullah, Un Maddus, Abdul Haris Mustari, Mochamad Indrawan

## Abstract

The Peleng tarsier (*Tarsius pelengensis*) is a practically unknown prosimian, with a very narrow range limited to Banggai island-group, Central Sulawesi, Indonesia. It was classified as “Endangered” by IUCN in 2017. Detailed demographic and distribution information about *T. pelengensis* in the wild is not available even though they are crucial to setting up conservation priorities and strategies. The aim of this study was to analyze distribution, and population of *T. pelengensis* across the island group. Surveys were conducted over approximately five months in 2017 and 2018 across Peleng and the neighboring islands of Banggai, Labobo, and Bangkurung. It is now established that tarsiers occur on two of the major islands, namely Peleng and Banggai Island proper. Average density in Peleng and Banggai islands were estimated to be 247 individuals/km^2^, and this roughly fall within the broad ranges of tarsier densities in Sulawesi and offshore islands. In stark contrast to the only previously available supposition of distribution and conservation status (IUCN 2017), Tarsiers were found in nearly all elevations (0-937 m above sea level) and all habitats in Peleng island. There is also an entire newly discovered population in Banggai Island proper, which similarly distributed in varied habitats. Therefore, the conservation status of Peleng tarsier needs to be re-evaluated.

## Introduction

Tarsiers are spread across Southeast Asia, namely the Philippines, Malaysia, and Indonesia. There are currently 14 species of tarsiers in three genera: *Cephalopachus* in Kalimantan/ Borneo and Sumatera, *Tarsius* in Sulawesi and *Carlito* in Philippines [5] Of the 14 tarsier species, 13 are found in Indonesia and 12 in Sulawesi, including two new species. (Table 1).

**Tabel 1.**
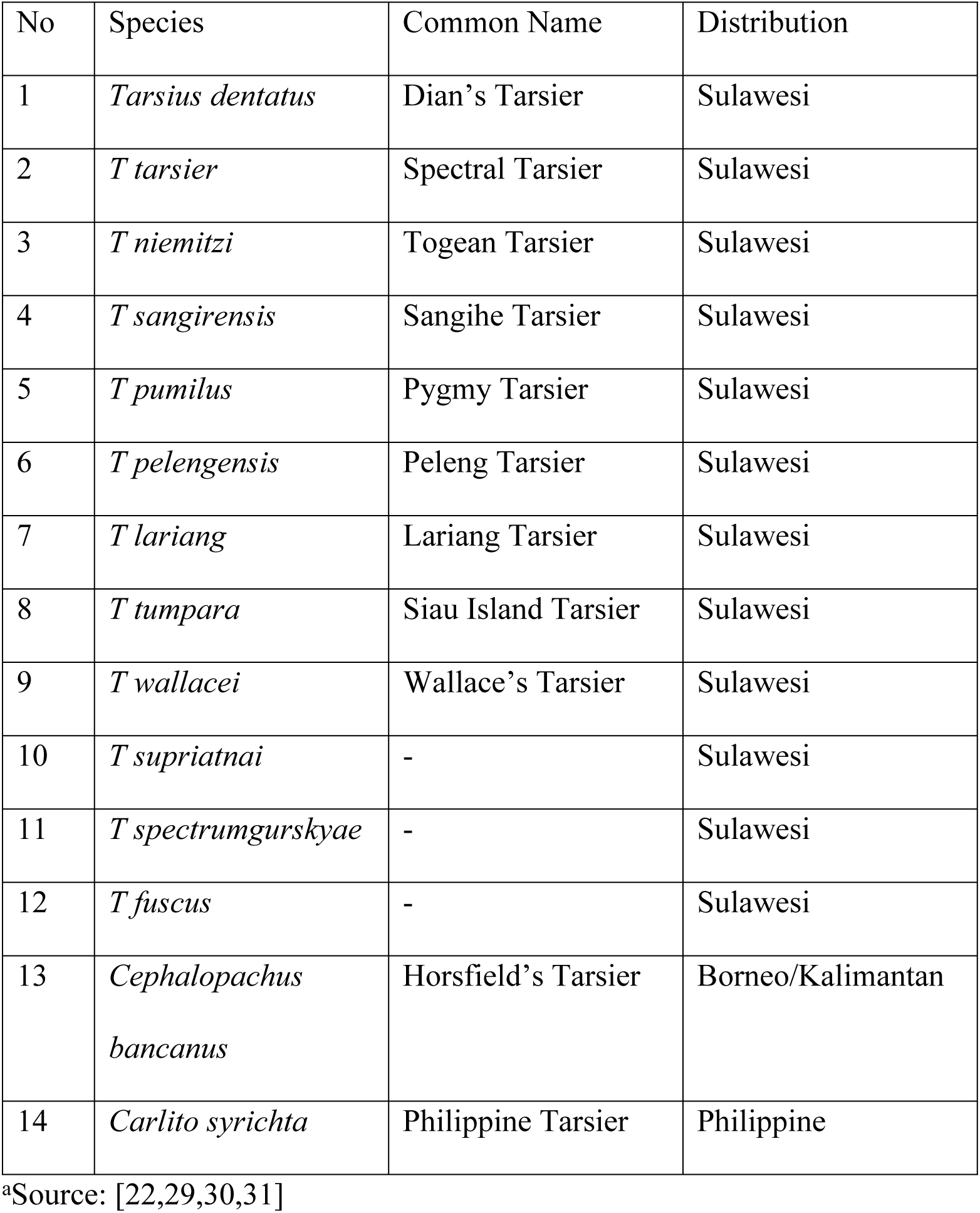
Tarsier species in Indonesia and Philippines

The Peleng tarsier *Tarsius pelengensis* is found only on Peleng Island, off the isthmus of Central Sulawesi [1,22,10,5]. Peleng is the largest island of the Banggai Island group, which also included Banggai, Labobo, and Bangkurung. Prior to this study, it was not known whether *Tarsius pelengensis* existed on the surrounding islands.

Despite its “Endangered” status [10,14] the ecology of Peleng tarsier is practically unknown. Detailed demographic and distribution information about *T. pelengensis* in the wild is not available but collecting demographic data is important to establishing the conservation priorities and strategies.

This study aims therefore to analyze the distribution, population, density and habitat use of Peleng tarsier also to re-evaluate its conservation status.

## Materials and methods

### Study site and subjects

The Banggai archipelago (Peleng Island, Banggai Island, Labobo Island, and Bangkurung Island) is vegetated by a mosaic of cultivation and scrubland with remnant natural ecosystems comprised of dryland tropical moist forests, swamp forests, mangroves, beach forests (Table 2).

**Tabel 2.** Comparison between area and forest cover in study site

| Island | Size (Km<sup>2</sup>) | Forest cover remaining<br>(Km<sup>2</sup>) |
| --- | --- | --- |
| Peleng Island | 2340 | 1172,9 |
| Banggai Island | 280 | 165.7 |
| Labobo Island | 73 | 55.8 |
| Bangkurung Island | 118 | 109,9 |

<sup>a</sup>extent estimated by the authors from published sources namely Republic of Indonesia Geospatial Information Agency 2018 and Global Forest Watch 2018

A previous study [12] highlighted that Peleng and surrounding islands had multiple endemic animals with threatened or near-threatened conservation status. Local wildlife population size was reduced by habitat degradation, predators, and trapping [12] which is even happening until today

This field study was conducted on the four main islands in the Banggai archipelago (Peleng, Banggai, Labobo, and Bangkurung) (Fig 1).

**Fig 1.**
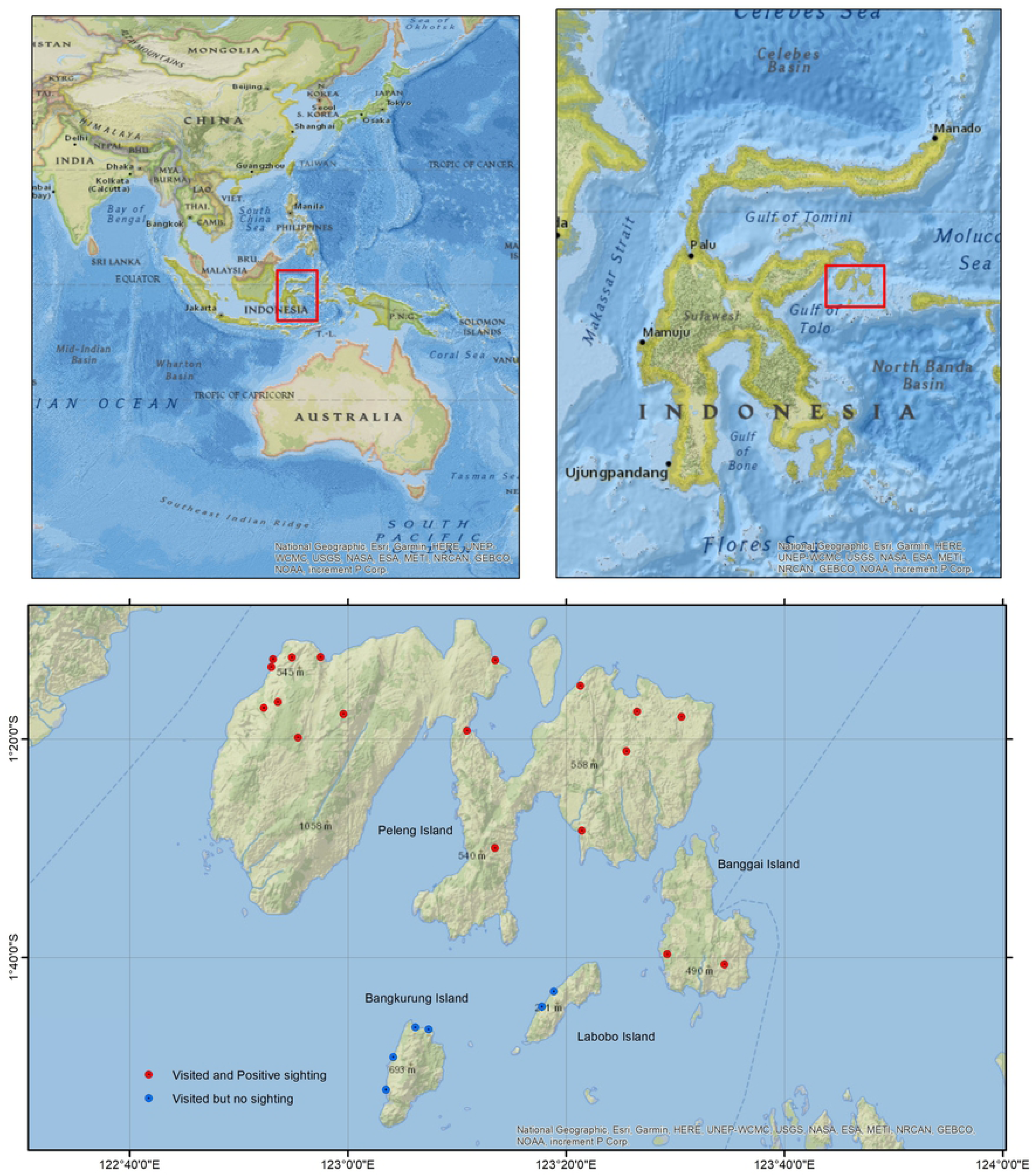
Map of the study area.

### Tarsier natural history

In the field, males and females appeared to exhibit dimorphism, with males relatively larger than females. Also, the males’ ears relatively longer than females’ and are pinkish near the tip pinkish. The ear tip of females was grayish.

Provisional natural history observations were continuously made by Un Maddus, whom born and raised around the forest margins, and a former hunter. UM regularly watched tarsiers since 2012 to this day and has familiarized himself with a singular female tarsier (“Simanis”, her being the only tarsier whom would raise her ears and turned her head when UM whispered her name from her vicinity). During the last 8 years, “Simanis” was regularly visited and watched by UM (without any attempt to provide food). The Tarsiers breed approximately once a year (usually at the beginning rainy season), and they produce a single infant whom began to stop nursing in about a 30 – 40 day’s time (UM, observations 2017 - 2020). “Simanis” was regularly encountered to this year when she gave birth to her 7^th^ offspring.

### Tarsier behavior ecology

In our field observation, tarsiers travel either alone or in groups of 2 to 7, whereas as many as 9 individuals have been encountered in one sleeping site. The group size of 2-7 individuals, was almost similar to size of groups of Dian’s Tarsier [19-21]. Groups were observed to use 1 - 3 sleeping trees in alternate days, whereas use of as many as 5 alternate sleeping trees have also been observed, possibly due to human disturbance. A similar behavior was noted in mainland Sulawesi, the tarsier frequently retreat to alternate shelter when disturbed [24]. Based on our preliminary study they foraged from 3-10 meters above ground (Un Maddus unpublished data)

Cursory observations of the tarsier in Peleng suggested that male and female ranged over areas of 10 ha and 8 ha, respectively. In mainland Sulawesi, the male tarsier moves further than the females [8] With the western tarsier *Tarsius bancanus*, [3] suggested that the male range was 11.25 ha while the female home range about 4.5 ha.

This species performs vocal duets at dawn before going to their sleeping site and infrequently at dusk. Tarsier species are known to be regularly vocalizing, and had dueted calls. In comparison, *Carlito* call less and *Cepalopachus* even less, and none of them were known to duet [5].

### Study methods

A 3-month (Dec. 2016 – March 2017) preliminary study focusing on Tarsier status and distribution was conducted by UM on Peleng Island and this yielded additional information about tarsier natural history. Tarsiers varied in their level of habituation. The tarsier in Banggai responded to playback calls of the tarsier from Peleng, and for this reason we are treating them as same taxonomic unit. Nevertheless, confirmation by genetic study is needed.

### Tarsier demographic surveys

Data were collected in April 7 – 23, 2017, July 18, 2017 to September 18, 2017 and March 19, 2018 to April 19, 2018 in Peleng Island and Banggai, Labobo, and Bangkurung from April 20 until May 19, 2018 for a total of 225 hours of contact. The population was sampled using transect method using the Distance sampling [2]. Transects were walked from 04:00 – 07:00, and 17:00 – 20:00. The transects were established using stratified random sampling. Twenty five transects were walked in Peleng, In Banggai, Labobo, and Bangkurung Islands, transects were walked 3-6 times.

The length of transect used was 800 m to 3,500 m, with the total length 74.1 km. The total area of the observation was about 10.01 km^2^. In location of study, determination of starting point of transect was random and stratified sampling was conducted over the 20 villages of Peleng, Banggai, Labobo, and Bangkurung Islands. The location of the detected tarsier groups was marked with GPS receiver (Table 3).

**Table 3.**
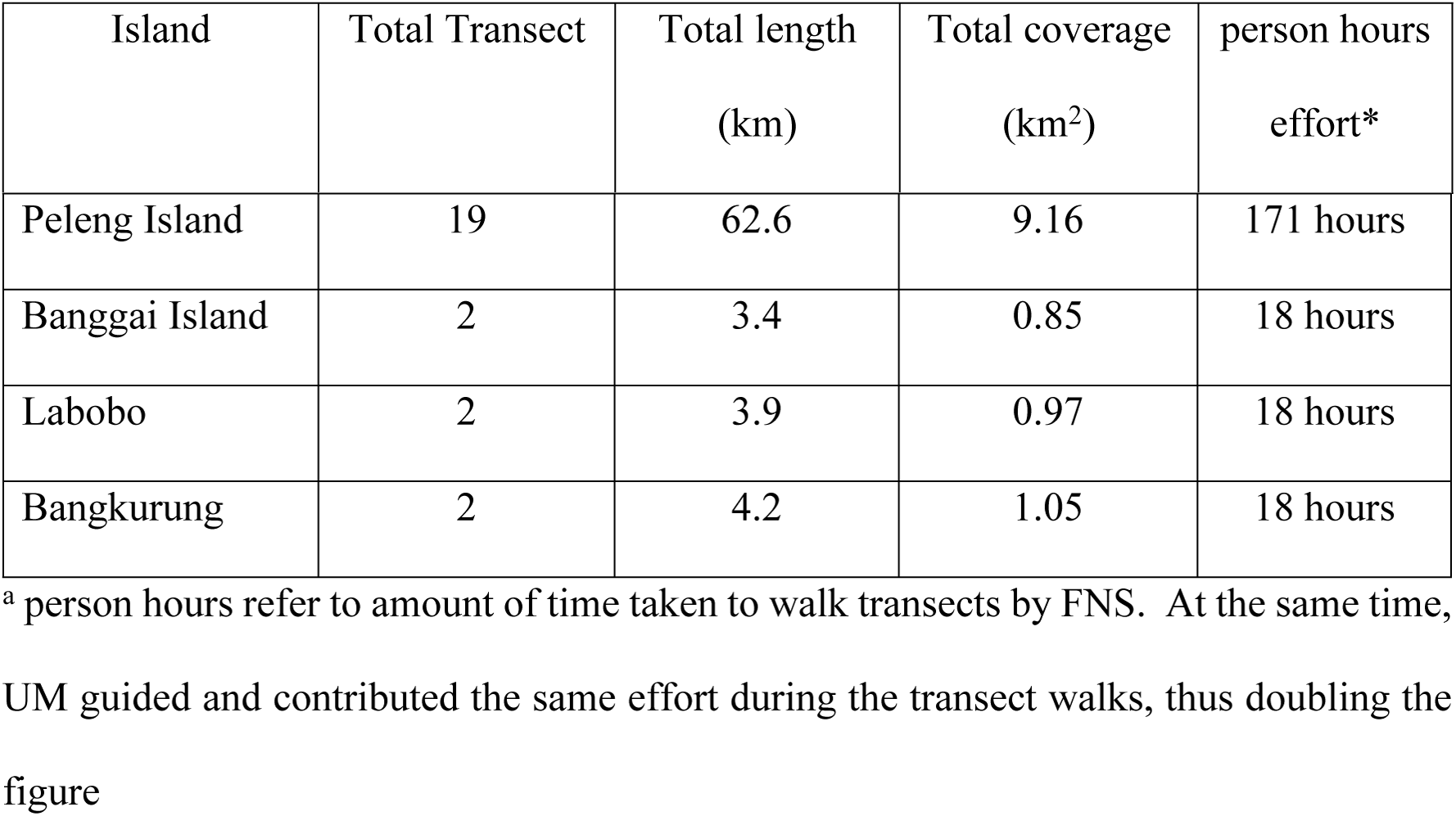
Transects used in tarsier surveys

Each group and individual encountered on transects were tallied which, yielded encounter rates and densities using software Distance 7.3 [2,32], using CDS (*Conventional Distance Sampling*). The model’s choice for the detection function was based on the smallest AIC (*Akaike’s Information Criterion*) value and the value of ΔAIC = 0 [32]. Hazard rate with simple polynomial was selected as primary function.

### Comparisons to other demographic surveys

Our method differed with Gursky [7] and Merker and Mühlenberg [19], whom used, respectively, fixed point count to estimate spectral tarsier densities and quadrat census distance to density conversion to estimate Dian’s tarsier densities

Gursky’s [7] modified form of the fixed-point count and quadrat census method computed total number of groups of *Tarsius spectrum* present in each hectare and this allows an estimate of the density for the sampled area. The plots were chosen randomly within the 1 km^2^ using random block design [7].

In Merker and Mühlenberg’s use of distance to density conversion, the number and spacing of tarsier family sleeping trees were considered to be a measure of habitat quality. Once all the sleeping trees in each habitat were known, the distance to their three nearest neighbor group were measured. The subsequent mapping of all sleeping sites resulted in estimates of population densities. [18, 19]. Among these studies none of the methodology has been standardize including this study. However, this study quite detailed.

There are assumptions in using distance sampling. objects are detected with certainty, object do not move, and measurements are exact [32]. Distance sampling requires large number of transect and randomized locations of transects. If the assumptions are not fulfilled biases will happen. In our case, the most likely source of bias is caused by failure in detection before moving. There is also the risk that the transects in Banggai, Labobo, and Bangkurung (but not Peleng) islands were too few.

### Habitat description work

We also conducted vegetation survey to understand the characteristic of the tarsier habitat. We used plotless sampling with point centered quarter method [34]. The observation points were placed on each transect where tarsiers were encountered and observed. A total of 50 points of vegetation observation were sampled. Herbarium specimens were prepared in the field, for a later identification by Mr. Ismail A Rahman, a botanical expert from Herbarium Bogoriense, Bogor, Indonesia. The data collected was analyzed to find the densities, and the importance values of the species.

## Results

This study established that Peleng tarsiers is geographically distributed in Peleng and Banggai island proper. As it was not found in Labobo and Bangkarung Island, the species might be practically absent in these two islands

Tarsiers were encountered in practically every habitat including secondary forests and even in the home gardens in villages such as Tatendeng, Leme Leme Darat, Okulo Potil, Kokolomboi, and Komba komba in Peleng.

The transects in Peleng Island and Banggai Island, respectively yielded direct sightings of 172 individuals and 13 individuals found during the sampling prosess. The estimated densities of Peleng tarsier was 247 Individuals/Km^2^ with interval from 157 - 391 individual/Km^2^. Coefficients of variation of the density was 22.57 % and transect effective width was 5.0224 m.

The results of Distance software analysis which indicated average number of group members ranged from 1.7 to 2.01 individuals. Coefficients of variation of cluster size was 6.22%. However, that result was below the direct observations (2 – 7 individuals).

Our habitat study indicated the occurrence of distinct plant concorcia corresponding to altitudinal variation. The less disturbed forest tends to occur above 600 m and as high as 800 m.a.s.l. Vegetation analysis of the primary and secondary forest habitats yielded 69 species of plants from 36 families (Table 4).

**Table 4.** Important plants of tarsier habitat, across the altitudinal gradient in Peleng Island

| Local name | Family | Scientific name | Density<br>(individual/Ha) | Importance<br>Value (IV)<br>(%) |
| --- | --- | --- | --- | --- |
| **Altitudinal range 0-200 m asl (predominantly secondary forest)** |  |  |  |  |
| Kayu<br>tombos | Myrtaceae | *Syzygium glaucum*<br>(King) Chantaran.<br>& J. Prain | 248 | 75 |
| Bitaul | Clusiaceae | *Calophyllum*<br>*soulattri* Burm.f. | 83 | 21,7 |
| Tambade | Sapindaceae | *Mischocarpus*<br>*sundaicus* Blume | 124 | 18,6 |
| **Altitudinal 200-400 m asl (predominantly secondary forest)** |  |  |  |  |
| Sosom | Burseraceae | *Canarium*<br>*asperum* Benth. | 756 | 38 |
| Pilapuk | Elaeocarpaceae | *Elaeocarpus sp* | 378 | 36 |
| Tambade | Sapindaceae | *Mischocarpus*<br>*sundaicus* Blume | 504 | 28 |
| **Altitudinal range 400-600 mdpl (predominantly secondary forest)** |  |  |  |  |
| Osa | Fagaceae | *Castanopsis*<br>*acuminatissima*<br>(Blume) A. DC. | 247 | 52.4 |
| Onik | Dipterocarpaceae | *Shorea selanica*<br>(Lam.) Blume | 141 | 36 |
| Suloi | Fagaceae | *Lithocarpus*<br>*celebicus* (Miq.)<br>Rehder | 141 | 29,6 |
| **Altitudinal range 600-800 m asl (predominantly primary forest)** |  |  |  |  |
| Osa | Fagaceae | *Castanopsis*<br>*acuminatissima*<br>(Blume) A. DC. | 293 | 41,4 |
| Kanar | Myrtaceae | *Syzygium*<br>*fastigiatum*<br>(Blume) Merr. &<br>L.M. Perry | 586 | 36,3 |
| Takios | Lauraceae | *Cryptocarya*<br>*glauc*a Merr. | 74 | 33,1 |
| **Altitudinal range 800-1000 m asl (predominantly primary forest)** |  |  |  |  |
| Onik | Dipterocarpaceae | *Shorea selanica*<br>(Lam.) Blume | 33 | 34,8 |
| Suloi | Fagaceae | *Lithocarpus*<br>*celebicus* (Miq.)<br>Rehder | 66 | 31,2 |
| Kanar | Myrtaceae | *Syzygium*<br>*fastigiatum* | 50 | 29 |
|  |  | (Blume) Merr. &<br>L.M. Perry |  |  |

## Discussion

Encounter rates in Peleng island was 1.34 tarsier/Km and in Banggai island was 2.66 tarsier/Km. The latter estimate however may be confounded by smaller sample size. For instance, the Distance generated Coefficient of Variation from the encounter rate in Banggai island turned to be 85.3 percent. Nevertheless, the frequency of encounters in Banggai island was 3-9 individuals per km.

At any rate, since Banggai Islands has a modest forest coverage (Table 2), the figure seemed to confirm abilities of Peleng Tarsier to colonize degraded habitats thus again proving it is much less threatened than previously believed.

The calculated Peleng tarsier density in Peleng and Banggai Island of 247 individuals / km^2^ was relatively high (Table 5). Our tarsier detection rate in the field was 2-5 groups/km, or 2-15 individuals/km. This relative abundance range appeared to be consistent with Density-generated result.

**Table 5.**
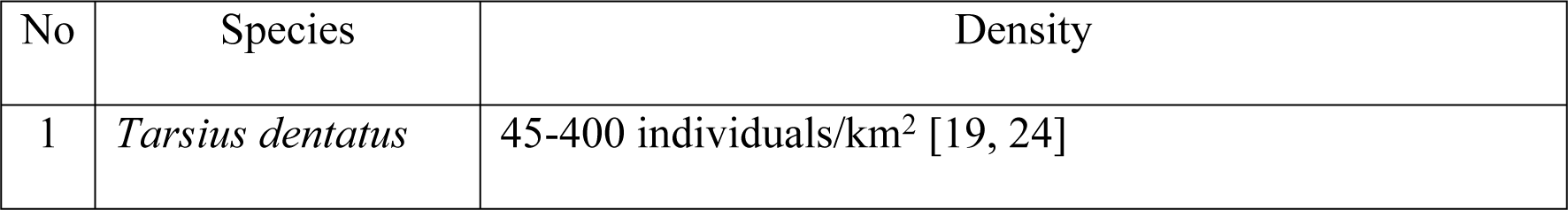
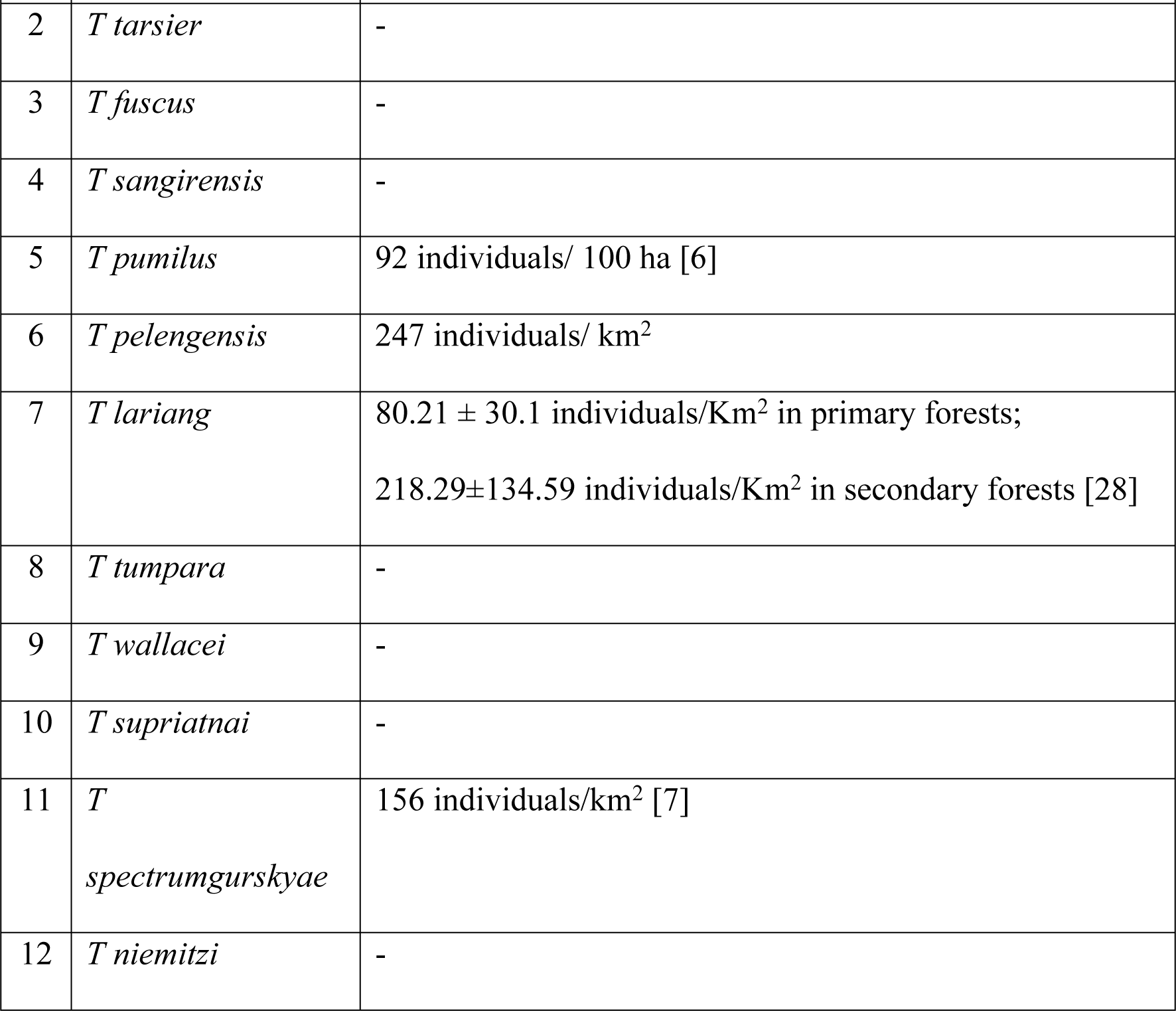
Estimated densities of different tarsier species

| No | Species | Density |
| --- | --- | --- |
| 1 | *Tarsius dentatus* | 45-400 individuals/km<sup>2</sup> [19, 24] |
| 2 | *T tarsier* | - |
| 3 | *T fuscus* | - |
| 4 | *T sangirensis* | - |
| 5 | *T pumilus* | 92 individuals/ 100 ha [6] |
| 6 | *T pelengensis* | 247 individuals/ km<sup>2</sup> |
| 7 | *T lariang* | 80.21 ± 30.1 individuals/Km<sup>2</sup> in primary forests;<br>218.29±134.59 individuals/Km<sup>2</sup> in secondary forests [28] |
| 8 | *T tumpara* | - |
| 9 | *T wallacei* | - |
| 10 | *T supriatnai* | - |
| 11 | *T spectrumgurskyae* | 156 individuals/km<sup>2</sup> [7] |
| 12 | *T niemitzi* | - |

The detection curve showed that only 28.1% of whole population were detected by the transect walks. This might happen because the tarsier moves before being recorded and detected.

For now, we cannot confirm that we had overestimated Peleng tarsier density. Detection probability this study was 28.1%. However when we truncated the data from 5% (outliers) we get 49,3% detection probability.

We are fairly confident of minimizing missing detection, for three reasons. Firstly, the maximum group size (7 individuals in a cluster) is on par with other tarsier species mentioned. Further the field studies is guided by the second author, whom knows tarsier all his 4 decades of life (a former hunter who is born and raised in the forest margins and even to the days of this write up continued to observe, monitor, and wrote admiring poems of the tarsier). The third author has monitored tarsier distribution weeks before and after the six months’ study period. Further, we found on 10 occasions that tarsier is attracted to moderately shoned flash lights, and this compensated or decreased the chances of missing detection.

### Group size

The group sizes of 2 – 7 individuals noted in the field (Peleng) were above the size generated from the Distance analysis (1.7 – 2 individuals in a group). Not all the individuals were detected during the observations and this might be a reason why made Distance-generated estimates had smaller group size compared to the direct encounters

In other tarsier species, the groups are showing various sizes, but generally in the range of Peleng tarsier’s, i.e. the largest group ranged 5 – 8. In the eastern tarsier, the group size of Pygmy tarsier was from 2 to 5 individuals [6] whereas Dian’s tarsier group live in small family of up to 7 individuals [21] and the western tarsier group size was reported 2 – 8 individuals [17].

### Factors potentially affecting tarsier density

Naturally, tarsier density is affected by various factors such as vegetation, predators, availability of food, and disturbance [19,35]. Each possibility will now be explored. In Peleng, tarsiers were more common at secondary forests than in primary forests. In general tarsiers have a widespread distribution and could live in various types of habitats such as secondary forests, primary forests, bamboo, shrubs and agricultural land [17]. With the western tarsier *T. bancanus*, it was also noted that compared to the large trees, sapling and poles were more attractive [3] Studies (e.g. [20]) have suggested that tarsiers would be more abundant in secondary forests with less disturbance than secondary forests with large disturbances. That tarsiers were more common in recently logged forests, shrubs, and bamboo or secondary forests with small-scaled disturbances is confirmed by a few studies already [25,23,4].

Forest fires and land clearing by burning happened regularly, and caused deforestation and degradation of habitat. Besides fisheries and services plantation was one of the livelihoods of the local community. The main agricultural commodities were tubers, and some agroforestry produces such as peanut (*Arachis hypogea*), cashew (*Anacardium occidentale*), fragrant nutmeg (*Myristica fragrans*), clove (*Syzygium aromaticum*). Intensive agriculture as a matter of course can reduce the species richness in the mosaic of the forested landscape [26]. Further, intensive land use may decrease mammalian density with the effect of small mammals greater than those of larger sized [33]. In Peleng, wildlife-threatening forest fires happened in high of dry season in 2015, within the administrative boundaries of Okulo Potil village (Yoris Yosukule pers. comm, 31 Oct 2017). For the local community, use of fire in agriculture was the easiest, quickest, and cheapest way to clear land (see also [11]). Again though, such an incident had not proven to be detrimental to the tarsier, at least in the short term.

Tarsiers preys on insects such as grasshopper (Orthoptera), fireflies (Coleoptera), and spiders (Aranae) (Alpian Maleso, pers comm, 24 Aug 2017). In mainland Sulawesi, tarsiers were known to eat insects from the orders of Aranae, Coleoptera, Isoptera, Homoptera, Hymenoptera, Lepidoptera, and Orthoptera [8]. The western tarsier *T bancanus*, was even known to feed on small snakes and lizards [16,4]. Foraging behavior and diet preferences of tarsiers were affected by season. In rainy season, their diet preferences may different in mainland Sulawesi. In the rainy season tarsiers will tend to hunt Orthoptera and Lepidoptera [8]. In the Sulawesi moist forests, insects is the main diet of most commonly found in secondary vegetation of forest edges [6].

It may be that for Peleng Island more insects were found in secondary. It is not known whether temperature is a limiting factor for altitudinal distributional factor of tarsier

In Sulawesi and the Philippines tarsiers was predated by snakes, lizards, owls, eagles, cats, civets [27], with *Python reticulatus* as potentially important predator [9]. Python, lizards, eagle, owl, civet also occurred in Peleng island and thus are potential tarsier predators. In one observation the rather common *Spilornis rufipectus* Sulawesi Serpent-eagle was stealthily watching a lone tarsier (Un Maddus, unpublished).

### Conservation status

The current status of the Peleng tarsier based on [14] was Endangered (EN) B1ab(iii) which means that the geographic distribution is smaller than 5000 km^2^, the density estimated show serious fragmented population or could be found only at one location and decreasing area or habitat quality [15]. Gursky [10] suggested that suitable habitat for tarsiers was only 10% of the island based on assumption of habitat extent but until our present research there has not been any proper field survey of the Peleng tarsier let alone detailed demographic study.

However, based on our study, the total population of the Peleng tarsier was estimated to be 2,270 individuals from 1,001 Ha. We estimated almost 537,225 individuals on Peleng island and Banggai island. This result suggests that *T. pelengensis* does not fit into any IUCN “threatened” criteria. Tarsiers have spread into a variety of natural ecosystems in Peleng and Banggai Island therefore they do not meet the Endangered criteria B1ab (iii). Hence, we propose that the Peleng tarsier to be reduce from Endangered (EN) to Near Threatened (NT).

The Peleng tarsiers are not hunted and found on both Peleng and Banggai islands. Increased habitat protection mainly by communities as part of longer-term local people-centered capacity building [13] has been effective strategy. As follow up to the fieldtrip series [13] the local community has independently conducted their own wildlife conservation outreach, and a growing number of community-conserved areas have been established in the villages of Kokolomboi/ Lemeleme Darat, Komba-komba, Kawalu/ Kautu, and Batong.

## Conclusions

Peleng tarsier (*T. pelengensis*) as it is currently known from the 2 neighboring islands of Peleng and Banggai, Central Sulawesi, Indonesia, was encouragingly found across highly varied habitat types, and was estimated to have a total density of 247 individuals/km^2^. Considering our 6 month’s of field data (in contrast rough assumption that lacked actual field data that formed bases on the previous “Endangered” classification) demonstrated clear abundance and adaptation to high levels of human disturbances in the two islands, we would like to suggest the tarsier to be declassified from Endangered status category, and move into “Near Threatened (NT)”.

## Acknowledgements

Our thanks go to the Banggai Island islanders and including the Togong Tanga community who supported our activities and shared their indigenous knowledge. Further support are given by the government of Banggai Is district, including Mssrs Kondrad Galala and Ferdy Salamat. For additional information on tarsier natural history we thank Ms. Lisa Nurfalah, Mssrs Labi Mopook and Alpian Maleso. Many thanks to Professor Sharon Gursky and Dr Stefan Merker. We thank Mssrs Jens-Ove Heckel, Roland Wirth, Noel Rowe for their kind facilitation and interesting discussions especially during the conception of the tarsier field study.

## Author contribution

Concept and writing of the manuscript: FN. Syahrullah, AH. Mustari and M. Indrawan Statistical analyses, maps and figures: FN. Syahrullah, AH. Mustari and M. Indrawan Data collection, provision of logistic, materials and site access: FN. Syahrullah, U. Maddus, and M. Indrawan

## References

1. Brandon-Jones D, Eudey AA, Geissmann T, Groves, CP, Melnick DJ, Morales JC, Stewart CB. Asian Primate Classification. International Journal of Primatology. 2003; 25(1): 97–164.

2. Buckland ST, Rexstad EA, Marques TA, Oedekoven CS. Distance Sampling: Methods and Applications. London: Springer; 2015

3. Crompton RH, Andau PM. Locomotion and habitat utilization in free ranging *Tarsius bancanus*: a preliminary report. Primates. 1986; 27(3): 337–355.

4. Fleagle JG. Primate Adaptation and Evolution. San Diego, US: Elsevier Inc; 2013

5. Groves C, Shekelle M. The genera and species of *Tarsiidae*. International Journal of Primatology. 2010; 31(6): 1071–1082.

6. Grow N, Gursky S, Duma Y. Altitude and forest edges influence the density and distribution of pygmy tarsier (*Tarsius pumilus*). American Journal of Primatology. 2013; 75: 464–477.

7. Gursky S. Conservation status of the spectral tarsier, *Tarsius spectrum*: population density and home range size. Folia Primatology. 1998; 69:191–203.

8. Gursky S. Effect of seasonality on the behavior of an insectivorous primate, *Tarsius spectrum*. International Journal of Primatology. 2000; 21(3): 477–495.

9. Gursky S. Predator mobbing in *Tarsius spectrum*. International Journal of Primatology. 2005; 26(1): 207–222.

10. Gursky S, Shekelle M, Nietsch A. The conservation status of Indonesia’s tarsier. In: Shekelle M, Maryanto I, Groves C, Schulze H, Fitch-Snyder H, editors. Primates of the oriental night. Bogor: LIPI Press; 2008. pp. 105–114

11. Harrison ME, Page SE, Limin SH. Global impact of Indonesia forest fires. Biologist. 2009; 56(3): 156–163.

12. Indrawan M, Fujita MS, Masala Y, Pesik L. Status and conservation of sula scrubfowl (*Megapodius bernsteinii* Schlegel 1866) in Banggai Islands, Sulawesi. Tropical Biodiversity. 1993; 1(2): 113–130.

13. Indrawan M, Garnett ST, Masala Y, Wirth R. Compromising for conservation: a protocol for developing sustainable conservation plans in biologically rich and monetarily impoverished communities. Pacific Conservation Biology. 2014; 20 (1): 3–7.

14. [IUCN] International Union for Conservation Nature. Peleng tarsier conservation status. 2008 June 30 [cited 17 March 2018]. In : Red List of Threatened Species [Internet]. Available from: https://www.iucnredlist.org/species/21494/9290015.

15. [IUCN] International Union for Conservation Nature. IUCN Red List Categories and criteria. Gland: IUCN; 2012.

16. Jablonski N, Crompton RH. Feeding behavior, mastication, and tooth wear in the western tarsier (*Tarsius bancanus*). International Journal of Primaology. 1994; 15(1): 29–59.

17. Mackinnon J, Mackinnon, K. The behavior of wild tarsier. International Journal of Primatology. 1980; 1(4): 361–379.

18. Merker S, Mühlenberg M. Traditional land-use and tarsiers – Human influences on population densities of *Tarsius dianae*. Folia Primatologica. 2000; 71: 426–428

19. Merker S, Yustian I, Mühlenberg, M. Losing ground but still doing well-*Tarsius dianae* in human-altered rainforest of Central Sulawesi, Indonesia. In: Gerold G, Fremerey M, Guhardja E, editors. Land Use, Nature Conservation and Stability of Rain Forest Margins in Southeast Asia. Heidelberg: Spinger; 2004. pp. 299–311.

20. Merker S, Yustian I, Mühlenberg M. Responding to forest degradation: altered habitat use by Dian’s tarsier *Tarsius dianae* in Sulawesi, Indonesia. Oryx. 2005; 39(2): 189–195.

21. Merker S. Habitat-Specific Ranging Patterns of Dian’s Tarsiers (*Tarsius dianae*) as Revealed by Radiotracking. American Journal of Primatology. 2006; 68: 111–125.

22. Merker S, Groves CP. *Tarsius lariang*: A new primate species form Western Central Sulawesi. International journal of primatology. 2006; 27(2): 465–485.

23. Merker S, Yustian I. Habitat use analysis of dian’s tarsiers (*Tarsier dianae*) in mixed species plantation in Sulawesi, Indonesia [Short communication]. Primates. 2007; 49: 161–164.

24. Merker S. The population ecology of dian’s tarsier. In: Gursky-Doyen S, Supriatna J. editors. Indonesian Primates. New York: Springer: 2010. pp. 371–382.

25. Niemitz C. (1979). Outline of the behavior of *Tarsius bancanus*. In: Doyle GA, Martin RD. editors. The study of prosimian behavior. London: Academic Press, Inc: 1979. pp. 631–660.

26. Phillips HRP, Newbold T, Purvis A. Land-use effects on local biodiversity in tropical forests vary between continents. Biodiversity Conservation. 2017; 26: 2251–2270.

27. Řeháková-Petrů, M, & Peške L. Predation on a wild philippine tarsier (*Tarsier syrichta*) [Short commuication]. Acta ethologica. 2012; 15: 217–220.

28. Rosyid A. Demography Parameter Estimation and Habitat Preference of Lariang Tarsier (*Tarsius lariang*) in Lore Lindu National Park Area. Journal of Natural Resources and Environmental Management. 2019; 9(1): 144–151.

29. Shekelle M, Groves C, Merker S, Supriatna J. Tarsius tumpara: a new tarsier species from Siau Island, North Sulawesi. Primate Conservation. 2008; (23): 55–64.

30. Shekelle M, Groves CP, Maryanto I, Mittermeier RA. Two new tarsier species (tarsiidae, primates) and the biogeography of Sulawesi, Indonesia. Primate Conservation. 2017; (31).

31. Shekelle M, Groves CP, Maryanto I, Mittermeier RA, Salim A, Springer MS. A new tarsier species from the Togean Islands of Central Sulawesi, Indonesia, with references to Wallacea and conservation on Sulawesi. Primate Conservation. 2019; (33)

32. Thomas L, Buckland ST, Rexstad EA, Laake JL, Strindberg S, Hedley SL. Distance software: design and analysis of distance sampling surveys for estimating population size. Journal of Applied Ecology. 2010; 47:5–14.

33. Wearn OR, Rowcliffe JM, Carbone C, Pfeifer M, Bernard H, Ewers RM. Mammalian species abundance across a gradient of tropical land-use intensity: a hierarchical multi-species modelling approach. Biological conservation. 2017; 212:162–171.

34. Wheater CP, Bell JR, Cook PA. Practical Field Ecology: A Project Guide. Chichester: Wiley Blackwell; 2011

35. Yustian I, Merker S, Supriatna J, Andayani N. Relative population density of *Tarsius dianae* in man-influenced habitats of Lore Lindu National Park, Central Sulawesi, Indonesia. Asian Primates Journal. 2008; 1(1):10–16.

